# Transcriptomic analyses of MYCN-regulated genes in anaplastic Wilms’ tumour cell lines reveals oncogenic pathways and potential therapeutic vulnerabilities

**DOI:** 10.1101/2021.01.11.426177

**Authors:** Marianna Szemes, Zsombor Melegh, Jacob Bellamy, Ji Hyun Park, Biyao Chen, Alexander Greenhough, Daniel Catchpoole, Karim Malik

**Affiliations:** Cancer Epigenetics Laboratory, School of Cellular and Molecular Medicine, University of Bristol, Bristol, UK.; Department of Cellular Pathology, Southmead Hospital, Bristol, UK.; Applied Sciences, University of the West of England. Bristol, UK.; The Kids Research Institute, The Children’s Hospital at Westmead, Westmead, New South Wales 2145, Australia.

**Keywords:** Wilms’, tumour, MYCN, REST, PRMT, TOMM20, RNA-seq

## Abstract

The *MYCN* proto-oncogene is deregulated in many cancers, most notably in neuroblastoma where *MYCN* gene amplification identifies a clinical subset with very poor prognosis. Gene expression and DNA analyses have also demonstrated over-expression of *MYCN* mRNA, as well as focal amplifications, copy number gains and presumptive change of function mutations of *MYCN* in Wilms’ tumours with poorer outcome, including tumours with diffuse anaplasia. Surpisingly, however, the expression and functions of the MYCN protein in Wilms’ tumours still remain obscure.

In this study, we assessed MYCN protein expression in primary Wilms’ tumours using immunohistochemistry of tissue microarrays. We found MYCN protein to be expressed in tumour blastemal cells, and absent in stromal and epithelial components. For functional studies, we used two anaplastic Wilms’ tumour cell-lines, WiT49 and 17.94, to study the biological and transcriptomic effects of MYCN depletion. We found that MYCN knockdown consistently led to growth suppression but not cell death. RNA sequencing identified 561 MYCN-regulated genes shared by WiT49 and 17.94 cell-lines. As expected, numerous cellular processes were downstream of MYCN. MYCN positively regulated the miRNA regulator and known Wilms’ tumour oncogene *LIN28B*, the genes encoding methylosome proteins PRMT1, PRMT5 and WDR77, and the mitochondrial translocase genes *TOMM20* and *TIMM50*. MYCN repressed genes included the developmental signalling receptor *ROBO1* and the stromal marker *COL1A1*.

Importantly, we found that MYCN also repressed the presumptive Wilms’ tumour suppressor gene *REST*, with MYCN knockdown resulting in increased REST protein and concomitant repression of REST target genes. Together, our study identifies regulatory axes that interact with MYCN, providing novel pathways for potential targeted therapeutics for poor prognosis Wilms’ tumour.

## 1. Introduction

Wilms’ tumour (WT) is the most common paediatric renal malignancy. WT can broadly be categorized as favourable histology (FHWT) or anaplastic. Whilst survival of FHWT patients after neoadjuvant therapy has improved overall survival, patients often relapse and experience extensive side-effects as a result of current therapies, with survivors remaining at elevated risk for death long after their diagnosis. Stage III-IV tumours, including anaplastic WT can have markedly worse prognosis, with a 4-year survival rate as low as ~50%. Thus there remains a critical requirement for more personalised, targeted therapies to prevent severe illness and death from WT (1,2).

The earliest genetic analyses of WT showed loss-of-function mutations in *WT1* (3,4), missense *TP53* mutations (5) and gain-of-function *CTNNB1* mutations resulting in activation of Wnt signalling (6). These mutations segregate with WT subtypes, for example *WT1* and *CTNNB1* mutations in stromal-predominant WT, and *TP53* in anaplastic WTs (5,7,8). More recent genome sequencing studies have found further mutations, including *MYCN, REST, SIX1/2, DROSHA* and *DICER* (7-9), reported in approximately half of all WTs. Inactivating mutations of *REST* were also independently reported, implicating it as a WT tumour suppressor gene in familial and non-familial WT (10).

Together with previous studies demonstrating *MYCN* gain and *FBXW7* loss associated with diffuse anaplasia and poorer outcome even in the absence of anaplasia, and focal amplifications of *MYCN* in anaplastic WTs (11,12), these whole-genome sequencing analyses suggest an oncogenic role for MYCN in WT. This is further supported by the fact that several groups documented *MYCN* mRNA over-expression in WTs, and its association with poor prognosis (13–15). Together this strongly implicates *MYCN* deregulation in Wilms’ tumorigenesis. However, despite MYCN being known to be important in proliferation of mesenchymal progenitor cells during nephrogenesis (16) and an established oncogenic transcription factor of developmental cancers such as medulloblastoma and neuroblastoma (NB) (17), virtually nothing is known about the biological activities of MYCN in WT, including protein expression patterns, downstream transcriptional targets, and possible pathways regulated.

In this study, we report the first analysis of MYCN protein in primary WTs. Furthermore, our functional analyses demonstrate that MYCN regulates proliferation of anaplastic WT cell-lines. RNA sequencing of these cell-lines after MYCN depletion identifies novel growth control pathways regulated by MYCN, including intersection with the function of the putative WT tumour suppressor gene, *REST* (RE1-Silencing Transcription factor).

## 2. Materials and Methods

### 2.1. Wilms’ tumour cell-lines, culture conditions and siRNA treatments

Wit49 (18) and 17.94 (19) anaplastic WT cell-lines were kind gifts from Prof. Herman Yeger and Dr. Keith Brown, respectively. The identity of both cell-lines was confirmed by short tandem repeat (STR) analysis. Wit49 cells were cultured at 37°C under 5% CO_2_, in Dulbecco’s modified Eagle’s medium supplemented with 15% fetal calf serum, 2 mmol/L L-glutamine, 0.1 mg/mL penicillin/streptomycin, 0.6%(v/v) β-mercaptoethanol and 1x insulin–transferrin–selenium, all purchased from Sigma. 17.94 cells were grown in Dulbecco’s modified Eagle’s medium supplemented with 10% fetal calf serum, L-glutamine and penicillin and streptomycin. Absence of Mycoplasma infection was confirmed by Mycoalert Mycoplasma Detection Kit (Lonza). Knock-down experiments were performed by using RNAiMAX reagent (Invitrogen) with 20 nM siRNA, according to manufacturer’s instructions. Oligonucleotide sequences are shown in Supplementary Table 1.

### 2.2. Cell cycle analysis and cell counting

Propidium-iodide labelling and fluorescence activated cell sorting (FACS) analysis was used to detect cell cycle phases. Floating and adherent cells were collected, washed with PBS, fixed with ice cold 70% (v/v) ethanol and treated with RNase A (Qiagen). After adding 50 µg/mL Propidium Iodide (Sigma), the samples were incubated at 37°C for 15 minutes, and analysed on Fluorescence Activated Cell Sorter LSRFortessaTM X-20 (BD Biosciences). About 15,000 events were collected for each replicate and data was analyzed by using FlowJo software. Cell counting was performed by using Countess automated cell counter (Invitrogen) and trypan blue staining.

### 2.3. Immunohistochemistry

Tissue microarrays, containing 33 pre-chemotherapy, Wilms’ tumor samples and fetal and adult kidney sections were stained by using a MYCN antibody (Proteintech, 10159-2-AP, Lot no: 18121). Immunohistochemistry staining was scored as positive or negative by a pathologist blinded to the specimens. All human tissues were acquired in compliance with the NSW Human Tissue and Anatomy Legislation Amendment Act 2003 (Australia). Ethics clearances 09/CHW/159 and LNR/14/SCHN/392 were approved by the Sydney Children’s Hospital Network Human Research Ethics Committee to construct TMAs and use clinical data, which was deidentified. Immunohistochemistry was performed with a Leica Microsystem Bond III automated machine by using the Bond Polymer Refine Detection Kit (Ref DS9800) followed by Bond Dab Enhancer (AR9432). The slides were dewaxed with Bond Dewax Solution (AR9222) and heat mediated antigen retrieval was performed using Bond Epitope Retrieval Solution for 20 minutes.

### 2.4. Protein Extraction and Western Blot

Cells were lysed in Radioimmunoprecipitation assay (RIPA) buffer and protein concentration was determined by using Micro BCA TM protein assay kit (Thermo Fisher). Protein extracts were loaded onto SDS poly-acrylamide gels and run in 1x Tris-glycine SDS buffer. After transfer onto PVDF membrane (Millipore) by a wet protocol (Bio-Rad), the membrane was blocked in 5% (w/v) skimmed milk, incubated with primary antibody solution at 4 °C overnight, and HRP labelled secondary antibody solution the next day. Proteins were visualised by using ECL reagents (Lumiglo, KPL) and X-ray films. The antibodies used are listed in Supplementary Table 2. Western blot image data were quantified by using ImageJ software (http://imagej.nih.gov/ij/). Target protein band density was normalized to the respective loading control and to the normalized intensity of the control sample.

### 2.5. RNA extraction, reverse transcription and qPCR

RNA was extracted by using the miRNeasy kit (QIAGEN), according to manufacturer’s instructions. RNAs were treated with on column DNase digestion using RNase-free DNase (Qiagen). RNA was transcribed with Superscript IV (Invitrogen) using a mixture of oligodT and random hexamer primers. Quantitative PCR was performed by using QuantiNova kit (Qiagen) on Mx3500P (Stratagene). The house-keeping gene TBP was used as a normalizing control. Relative gene expression was calculated using the ΔΔCt method – log2 fold changes between MYCN-depleted and control samples were calculated after normalization to TBP. Statistical significance of log2-transformed fold changes in gene expression was evaluated by using two-tailed, Student’s T-tests. The oligonucleotide primers used in this study are shown in Supplementary Table 1.

### 2.6. RNA-seq and bioinformatic analysis

Wit49 and 17.94 cells were treated with MYCN-targeting and control siRNAs for 48 hours, and were subsequently harvested in Qiazol (Qiagen). RNA was extracted by using the miRNeasy Kit (Qiagen). RNA concentration and quality were checked by using a Nanodrop spectrophotometer and Bioanalyzer (Agilent). Libraries were prepared from 200 ng RNA and were sequenced by using the paired-end option with 100 bp reads on BGIseq-500 (BGI Genomics). The reads were aligned to the human genome (hg38) by using STAR and the alignment files (BAM) files were further analysed in SeqMonk v1.47. (https://www.bioinformatics.babraham.ac.uk/projects/seqmonk/). Gene expression was quantified by using the Seqmonk RNA-seq analysis pipeline and differentially expressed genes (DEG) were identified by DESEQ2 (p<0.05), and a minimum shrunk fold difference threshold, a conservative corrected value of fold change taking confidence into account, of 1.2 was applied. RNA sequencing data is available from the European Nucleotide Archive (ENA) under the accession number ERP125499. Gene Signature Enrichment Analyses (GSEAs) were performed on preranked lists of log2-transformed, shrunk fold difference gene expression values (Broad Institute). Gene expression analysis of published Wilms’ tumour data sets and K means clustering were performed by using the R2 Genomics Analysis and Visualization Platform (http://r2.amc.nl).

### 2.7 Statistical analysis

Statistical analysis of quantitative PCR data was performed on log2-transformed fold change values, by using two-tailed Student’s T-tests. Gene Set Enrichment Analysis of RNA-seq data was evaluated based on Normalised Enrichment Score (NES) and False Discovery Rate (FDR), which was calculated based on permutation of genes with a rank score of log2 fold change expression over control. Differentially expressed genes (DEGs) in MYCN-depleted Wilms’ tumour cells, detected by using RNA-seq, were assessed using the statistical model implemented in DESEQ2. Significance of correlation between clusters of TARGET WT data and clinical correlates was assessed by using Chi-square test. Differential expression of genes and metagenes, which represent gene signatures, among groups of WT and fetal kidney tissue, was evaluated by using ANOVA.

## 3. Results & Discussion

### 3.1 MYCN protein is overexpressed in the blastemal component of Wilms’ tumour and promotes proliferation in Wilms’ tumour cells

Over-expression of *MYCN* mRNA has been reported in poor prognosis Wilms’ tumour (13–15,20) but the prevalence and pattern of MYCN protein expression in Wilms’ tumours has, to our knowledge, never been published. Therefore, we performed MYCN immunohistochemistry (IHC) on tissue microarrays (TMAs), containing 33 pre-chemotherapy WT sections (Figure 1). As a normal control, we used a section of a fetal kidney, derived from a 13-week old fetus, as well as an adult, healthy kidney. MYCN was detected in 14 tumours, exclusively in the blastemal component. It was localised mostly to the nucleus, but we also detected cytoplasmic staining, with three tumours displaying cytoplasmic staining only. In contrast, in normal fetal kidney, MYCN was detected solely in the distal tubules, but not in the blastema, while MYCN protein was completely absent from the adult kidney (Supplementary figure 1). Although the number of tumours precluded statistical analysis, our data demonstrates for the first time that MYCN protein is expressed in the blastemal cells of Wilms’ tumours.

**Figure 1.**
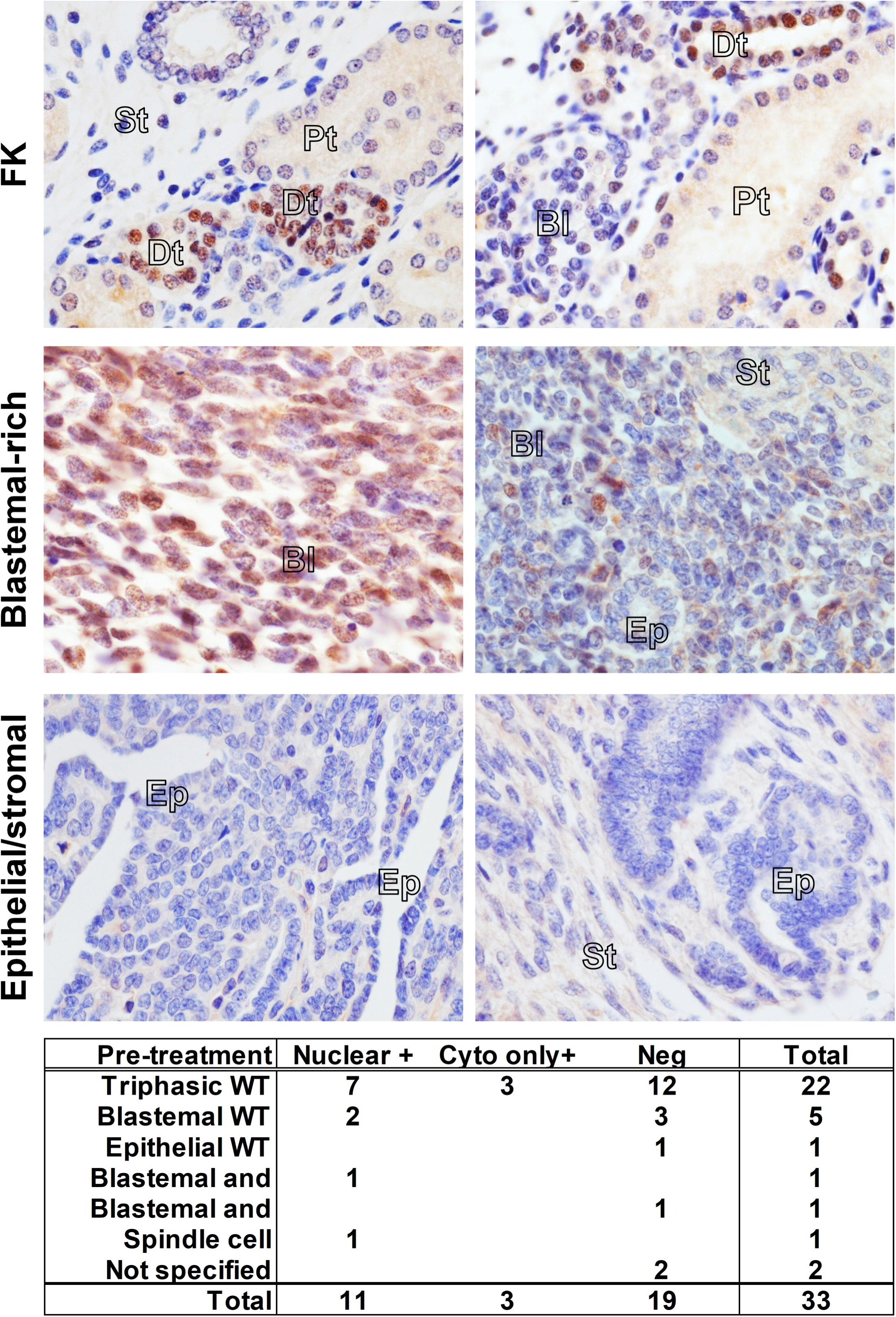
Immunohistochemistry in fetal kidney and Wilms’ tumours reveals a blastemal expression for MYCN protein in tumours. MYCN expression was detected in the distal tubules (Dt) in 13-weeks-old fetal kidney, while the blastema (Bl), stroma (St) and proximal tubules (Pt) were negative. Blastemal-rich WT showed positivity in the blastemal component. Epithelial and stromal structures did not show any expression of MYCN protein. A table summarizing positive MYCN staining in the nucleus or the cytoplasm only in WT, according to histology, is shown underneath.

To characterize the effect of MYCN on growth of Wilms’ tumour cells, we knocked it down in two anaplastic WT cell-lines, Wit49 and 17.94 (Figure 2). After 120 hours of MYCN depletion by two independent MYCN-targeting siRNAs, there was a substantial and significant reduction in live cell counts (p < 0.01). The number of dead cells did not increase, suggesting that the decrease in live cells was due to growth inhibition rather than increased cell death. To investigate the effect of MYCN depletion on the cell cycle, we performed a cell cycle analysis on Wit49 (Supplementary figure 2). MYCN knock-down by either siRNA led to a significant reduction of cells in S phase, while the proportion of cells in G2/M increased significantly (p < 0.01), indicative of a G2/M arrest. There was no increase of cells in the sub-G1 phase, in agreement with our previous observation of no increase in dead cell counts. These studies suggest that MYCN primarily exerts control over WT proliferative pathways as opposed to apoptosis and cell survival.

**Figure 2.**
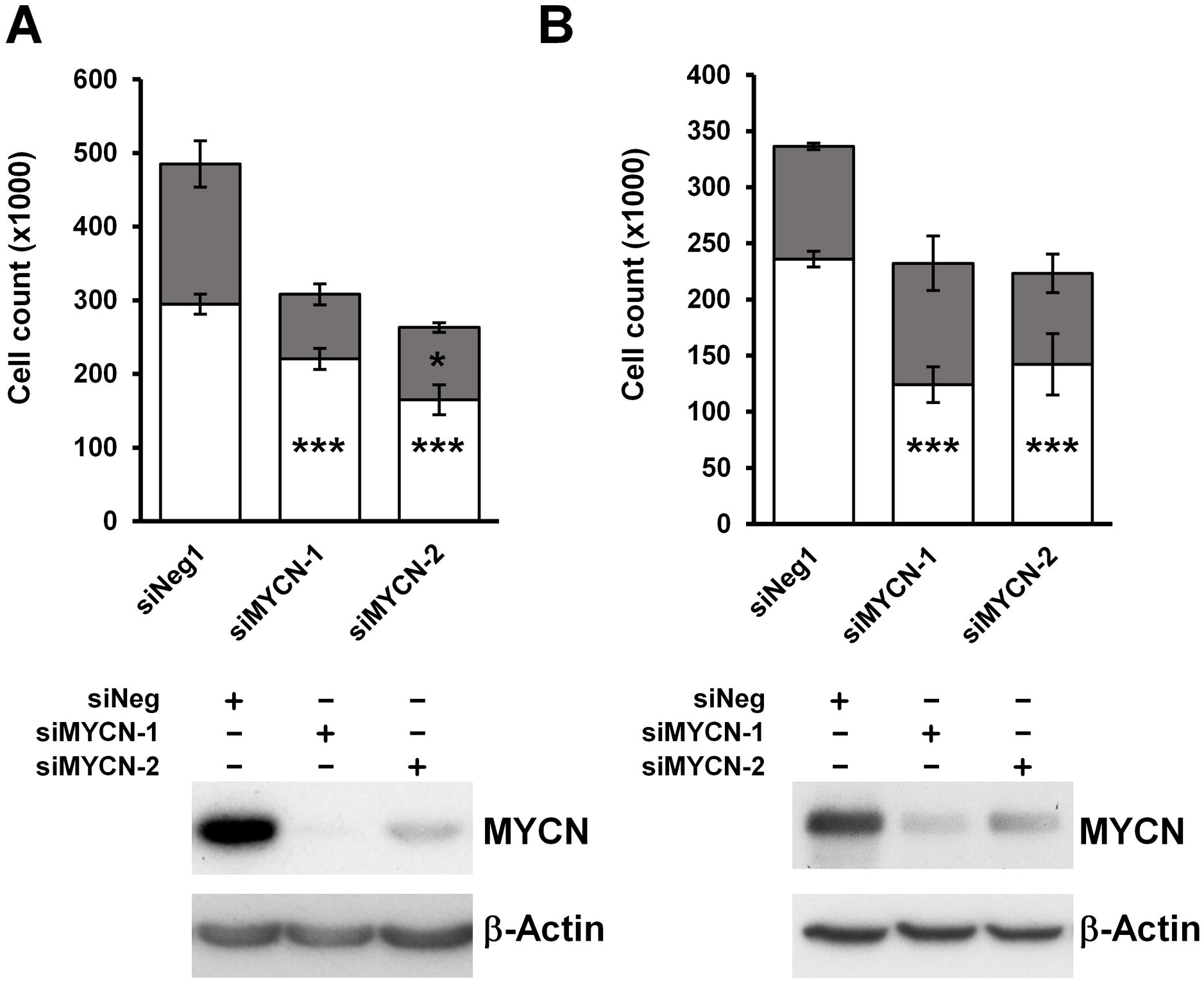
MYCN depletion leads to significant growth inhibition in anaplastic WT cell-lines. **(A)** MYCN depletion resulted in significant growth suppression after 120 hours treatment in Wit49, but not in an increase in dead cells, using two different siRNAs. (n = 3, *** p < 0.01, two-tailed T-tests). **(B)** MYCN knockdown also led to significant growth suppression in 17.94 cells after 120 hours (n = 3). Western blot was used to confirm knockdown of MYCN in both cell-lines.

### 3.2 MYCN-regulated gene signatures in Wilms’ tumour reveal downstream growth-regulatory pathways

To identify MYCN regulated genes in Wilms’ tumour, we performed RNA-seq of both anaplastic WT cell-lines following MYCN knockdown (Supplementary figure 3A). MYCN regulated genes were defined as differentially expressed genes (DEGs) that had a significant (p < 0.05) and substantial change (minimum ‘shrunk’ fold change, a corrected value based on confidence, of 1.2) in their expression levels, as assessed by using DESEQ2. We found 1060 upregulated genes in Wit49 and 396 in 17.94, with a highly significant overlap of 212 genes between the 2 cell-lines (p < 10^−10^) (Figure 3A). There were 349 downregulated genes shared by the two cell-lines, with 1086 and 699 identified in Wit49 and 17.94, respectively (p < 10^−10^). The shared 561 MYCN-regulated genes in WT are shown in Supplementary table 3. and Figure 3B. Known MYCN target genes, and genes related to WT or kidney development are highlighted on the heatmaps. A panel of the DEGs identified by RNA-seq after MYCN knockdown were validated in both cell-lines by qPCR. We confirmed downregulation of *LIN28B*, a MYCN target gene in neuroblastoma (21), both at the RNA (Figure 3C) and protein level (Supplementary figure 3B). LIN28B is an established regulator of nephrogenesis, promoting expansion of the progenitor pool, and a direct oncogenic driver in WT (22,23). LIN28B suppresses *let-7* miRNAs, but can also influence gene expression via other mechanisms, including regulation of translation (24). Therefore MYCN is likely to exert control at the post-transcriptional as well as the transcriptomic level.

**Figure 3.**
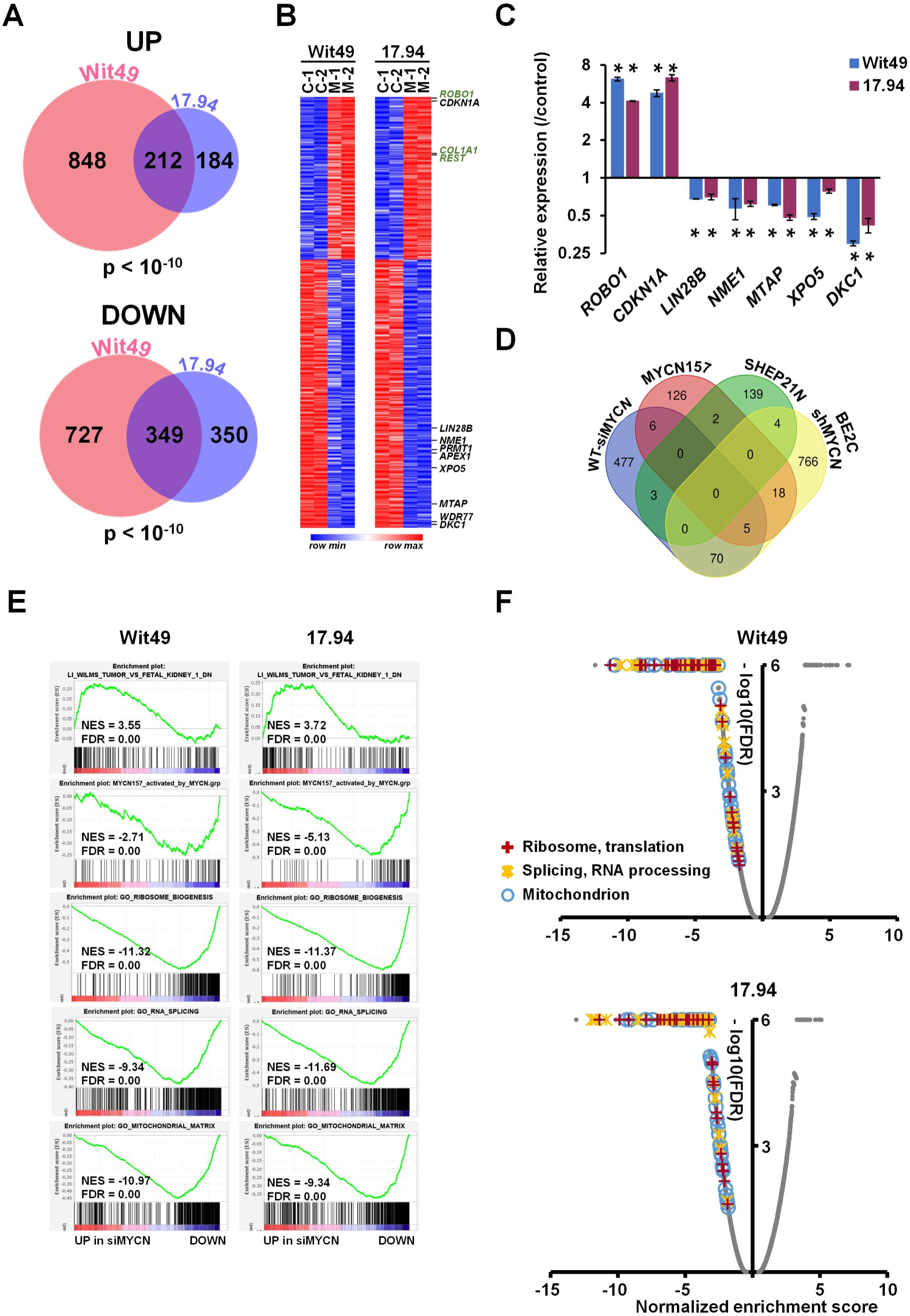
Genes and pathways identified by RNA-seq in MYCN-depleted anaplastic WT cells. **(A)** Venn diagrams showing highly significant overlaps between differentially expressed genes (DEGs) in Wit49 and 17.94 after MYCN depletion for 48 hours, for up- and downregulated genes, respectively. DEGs were determined as statistically significant (p < 0.05) and having a minimum shrunk fold change, a conservative corrected value, of 1.2, calculated by using DESEQ2. The probability values for shared genes is indicated. **(B)** Heatmap of shared 561 DEGs in Wit49 and 17.94. Examples of known MYCN target genes are shown in black, while genes associated with kidney development or Wilms’ tumour predisposition are highlighted in green. **(C)** Validation of identified, select MYCN-regulated genes in WT cells by qPCR, 48 hours after MYCN knockdown. A representative of three biological replicates is shown. Significance was calculated based on the biological replicates (* p < 0.05, T-tests). **(D)** Venn diagram showing overlaps of MYCN-regulated genes in WT, described in this study, with MYCN target genes identified in neuroblastoma. **(E)** GSEA plots showing upregulation of gene signatures repressed in WT as compared to FK, upon depletion of MYCN. MYCN-activated genes of the MYCN157 signature, identified in NB, were downregulated. Examples of GSEA of downregulated GO gene sets associated with ribosomal and mitochondrial function and splicing in MYCN depleted WT cells. **(F)** Volcano plots of Gene Set Enrichment Analysis on MYCN depletion transcriptomes in WT cells. Normalized enrichment scores and False discovery rates (FDR) were calculated using Gene Ontology gene categories indicated universal downregulation of gene sets linked to ribosomal and mitochondrial function as well as those associated with RNA processing and splicing. Scores of all gene sets are shown in grey, while those of statistically significant gene sets (FDR < 0.05) in the highlighted, functional categories are indicated according to the legend.

Amongst the novel genes, one of the biggest expression changes was the *ROBO1* gene, which increased over 4-fold after MYCN knockdown in both WT cell-lines (Figure 3C). *ROBO1* encodes the transmembrane Roundabout Guidance Receptor 1 involved in SLIT/ROBO signalling, a key developmental pathway (25). *Robo1* is a tumour suppressor gene in other cellular systems, with *Robo1* knockout mice predisposing to lung adenocarcinomas and lymphomas (26). Other studies have indicated that Slit/Robo signalling is required for normal kidney development (27), and *Robo1* expression increases in the pretubular aggregates compared to the metanephric mesenchyme, implying a role for Robo1 in early renal differentiation (28). We also note that *SLIT2* is frequently epigenetically silenced in WTs (29), further supporting a tumour suppressive role for SLIT/ROBO signalling in WT.

Gene set enrichment analysis indicated that MYCN repressed kidney differentiation and developmental genes (Supplementary figure 3C). To study how MYCN influences cell differentiation during nephrogenesis, we queried gene signatures characteristic of different cell populations in the fetal kidney, determined by Menon et al. by using single cell RNA-seq (30). We found that MYCN activated genes were overexpressed in proliferating cells, while gene signatures of both podocytes and stromal cells were repressed by MYCN, consistent with a role for MYCN in promoting growth and repressing differentiation (Supplementary Figure 3D). *COL1A1*, a marker gene for stromal cells, was upregulated more than two-fold in our RNA-seq data after MYCN depletion. Moreover, COL1A1 protein is documented to be significantly downregulated in anaplastic WT compared to favorable histology WT (31).

MYCN targets have been extensively studied in NB and several target lists were described. We compared our WT-specific MYCN targets to some identified by transcriptomic and chromatin immunoprecipitation analyses in NB, specifically (i) the MYCN157 signature, derived from IMR32 cell-line DEGs following MYCN knockdown, subsequently filtered by correlation with MYCN mRNA expression in primary NBs (32), (ii) DEGs from a MYCN-overexpressing isogenic model (33) and (iii) genes bound by MYCN and correlated with MYCN in the SEQC NB expression dataset (GSE49712) (34) (Figure 3D). We found surprisingly little overlap of our top DEGs with these NB-specific MYCN targets, emphasizing the importance of cellular context. To characterize the function of the newly identified MYCN regulated genes, we performed a Gene Ontology (GO) enrichment analysis (Supplementary table 4.). The most highly enriched component was the ‘RNA nuclear export complex’ (GO:0042565), including *XPO5* and *RAN*, responsible for the transport of pre-miRNAs from the nucleus, suggesting that MYCN may substantially alter the miRNA profile in the cytoplasm. Mutations in *DICER* and *DROSHA*, key enzymes in microRNA biogenesis, were previously shown to be involved in WT pathogenesis (8). The WT-specific MYCN regulated genes were also enriched in GO categories related to mitochondrial, ribosomal, spliceosomal and methylosome complexes and telomere maintenance, emphasising the control of several major cellular processes by MYCN.

To obtain an extended global overview of the transcriptional control of MYCN in WT, we performed Gene Set Enrichment Analyses (GSEA) with the DEGs from Wit49 and 17.94 cell-lines. GSEA revealed that MYCN knockdown led to activation of genes downregulated in WT relative to fetal kidney (Figure 3E) (35), and repression of genes elevated in WT relative to fetal kidney (Supplementary Figure 4A). Thus MYCN knockdown reverses, at least in part, the WT-specific signature.

**Figure 4.**
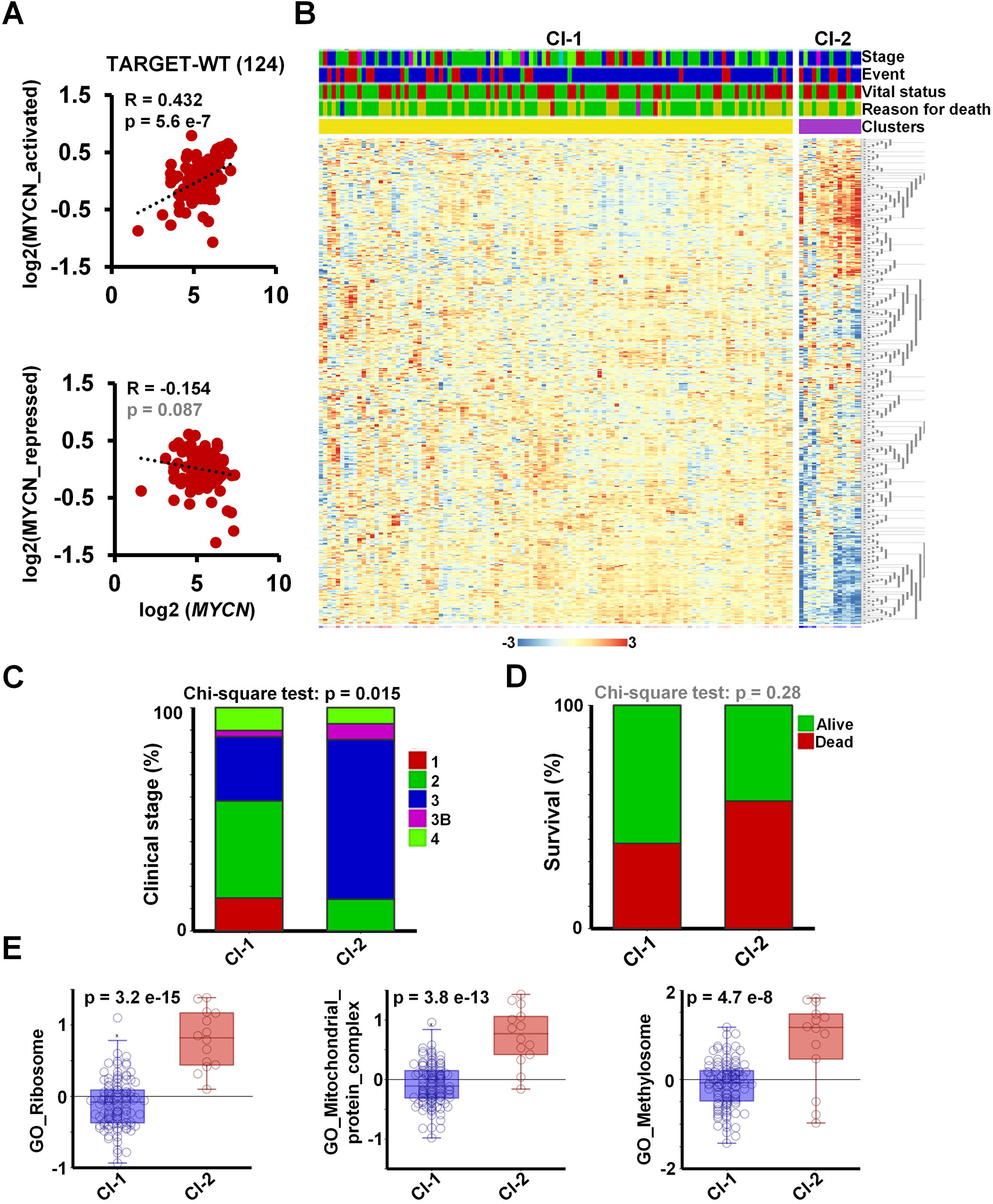
Meta-analysis of MYCN regulated genes in the WT TARGET data set identifies distinct patient clusters. **(A)** Overall expression of MYCN activated genes, identified as shared downregulated hits in MYCN depleted WT cells, showed a strong and significant positive correlation with MYCN expression in the TARGET-WT data set (SRP012006), containing transcriptomic data of 124 high-risk WT. Overall expression of MYCN repressed genes displayed a modest, inverse correlation that did not reach significance. **(B)** K means clustering (K = 2) performed on TARGET-WT data set based on the expression of 561 MYCN regulated genes in WT, identified in this study. Clinical information is shown a as coloured bars on the top: stage (red = 1, green = 2, blue = 3, magenta = 3b, light green = 4, yellow = 5_3B, turquoise = 5_4); event type (none = red, blue = relapse, green = progression); vital status (green = alive, red = dead) and reason for death (green = none, yellow = tumour, red = infection, blue = toxicity, magenta = tumour and toxicity). **(C)** Proportion of tumours with various clinical stages. Chi-square test showed that the association of clinical stages with clustering according to MYCN-regulated genes is significant. Stages represented by a single tumour were omitted. **(D)** More death occurred in patients with tumours in cluster 2, however this association did not reach significance. **(E)** Overall expression of ribosome genes, mitochondrial protein complexes and methylosome components was significantly higher in cluster 2 tumours as compared to those in cluster 1 (ANOVA).

MYCN activated genes in the MYCN157 signature were significantly downregulated, showing regulation of the NB-specific target genes in WT cells, despite the minimal overlap of our top DEGs (with the biggest or most significant changes) with top MYCN targets in NB. Similarly, several MYC target gene sets were downregulated in both cell-lines, indicating that MYCN drives canonical MYC target genes in WT (Supplementary figure 4B), like the curated MYC gene set in Hallmark (36). Further, genes encoding mitochondrial proteins that were recently shown to be regulated by MYCN (37) were also down-regulated, emphasizing the role of MYCN in activating mitochondrial function genes in WT. Genes encoding for proteins with roles in ‘RNA export from nucleus’ were down-regulated, which may affect pre-miRNA transport and the miRNA pool in the cytoplasm, while those participating in ‘Unfolded protein response’ were up, suggestive of endoplasmic reticulum stress in the MYCN depleted cells (Supplementary figure 4C).

Gene signatures related to ribosomal biogenesis, splicing, and mitochondria were all found to be down-regulated following MYCN knockdown, reinforcing the results of GO analysis with shared WT-specific MYCN-regulated genes (Figure 3E). In fact, all the Gene Ontology signatures related to to ribosomal function, RNA processing/splicing, and mitochondria were found to be profoundly downregulated in both MYCN depleted cell lines, suggesting a strong activation of these gene expression programmes by MYCN in WT (Figure 3F).

Ribosomal biogenesis was reported to be upregulated in *MYCN*-amplified neuroblastoma too, and inhibitors of RNA polymerase I (which transcribes ribosomal RNA genes) suppressed MYCN expression and promoted apoptosis of MNA NB cells, both *in vitro* and *in vivo* (38). Deregulation of splicing (39) and metabolic reprogramming by MYCN (40) were also observed in neuroblastoma, consistent with our transcriptomic analysis in WT.

To investigate the expression of *MYCN* and its target genes in a large set of primary Wilms’ tumour tissue, we analysed the publicly available WT RNA-seq data set, TARGET-WT (SRP012006), containing expression data for 124 high-risk tumours. We found a strong and highly significant, positive correlation between the expression of *MYCN* and its activated target genes in WT (R = 0.43, p = 5.6 × 10^−7^), suggesting a regulatory link *in vivo* (Figure 4A). The correlation between *MYCN* and the newly identified repressed genes in the tumours is less pronounced and does not reach significance (R = − 0.15, p = 0.087), suggesting that other regulators might also be involved in repression. For example, *LIN28B* was reported to be activated via chromosomal translocation and amplification in WT, independent of MYCN (41) and *REST* can be inactivated via mutations. To identify the tumours with a MYCN-regulated signature and study their clinical correlates, we clustered the WT transcriptomic data set according to the expression of the shared 561 MYCN target genes (Figure 4B). Two clusters were identified: a larger group (cluster 1) with a mostly uniform expression of target genes and a smaller group of 14 tumours (cluster 2), displaying up-regulated and down-regulated subsets of our MYCN target genes. The tumours in cluster 2 were generally higher stages, with a significant difference in distribution (Chi squared test p = 0.015, Figure 4C). Cluster 1 contained all the stage 1 tumours, and the proportion of stage 2 ones was higher as well (43% vs. 14%). In contrast, there was a higher proportion of stage 3 and 3B tumours in cluster 2: 71% vs. 28% and 7% vs. 3%, respectively. Patients with tumours in the MYCN-regulated group (cluster 2) had lower survival (42.9% vs. 61.8%), although this trend did not reach significance probably because of the relatively low number of patient samples in cluster 2 (Figure 4D).

To evaluate the expression of functional gene sets in primary Wilms’ tumours, we assessed the overall expression of GO gene categories highlighted by our RNA-seq analysis, represented as metagenes, in the newly defined clusters in TARGET-WT (Figure 4E). Ribosomal genes had a significantly and substantially higher expression in cluster 2 than cluster 1 (p = 3.2 × 10^−15^). Genes coding for mitochondrial protein complexes (p = 3.8 × 10^−13^) and methylosome components (p = 4.7 × 10^−8^) were similarly significantly upregulated in cluster 2, reinforcing our data in WT cell-lines. Together these analyses show that the DEGs we identified in Wit49 and 17.94 cell-lines correlate with MYCN in primary WTs, and that pathways identified by our *in vitro* analyses are aberrant in a subset of higher stage WTs.

### 3.3 MYCN upregulates key mitochondrial transporter gene TOMM20, overexpressed in blastemal and relapsed WT

Gene set enrichment analysis revealed a profound downregulation of genes encoding for mitochondrial complexes upon MYCN depletion, in both anaplastic WT cell-lines (Figure 5A). The most downregulated genes included *TOMM20* (Figure 5B), encoding a member of the translocase of the outer membrane (TOM) complex, responsible for the import of newly synthesized mitochondrial proteins from the cytosol. The TOM complex works in close co-operation with the translocase of the inner membrane (TIM) complex (42), a key member of which, *TIMM50*, was also downregulated with MYCN depletion. We confirmed the downregulation of these mitochondrial protein genes together with *PDK1* by qPCR (Figure 5C); PDK1 (Pyruvate Dehydrogenase Kinase 1) is a gatekeeper of the Warburg effect and frequently overexpressed in cancer (43). Downregulation of TOMM20 was also confirmed at the protein level in both anaplastic WT lines (Figure 5D). In the TARGET-WT RNA-seq dataset, we observed a significant, positive correlation between the expression of *MYCN* and *TOMM20*, consistent with our findings (Figure 5E). *TOMM20* was inferred to be a MYCN target gene in neuroblastoma due to MYCN binding to its promoter and its mRNA positively correlating with MYCN (34). However, our data at both protein and RNA levels demonstrate for the first time that TOMM20 is directly regulated by MYCN. In two other, publicly available transcriptomic data sets by Perlman et al. (44,45), *TOMM20* was found to be significantly overexpressed in relapsed WT vs. non-relapsed (Figure 5F) and in blastemal tumours relative to other tumours (Figure 5G). *TOMM20* over-expression is associated with poor prognosis in other cancers such as colorectal cancer and chondrosarcoma (46,47), and TOMM20 knockdown in colorectal cancer cells led to increased mitochondrial damage, significantly reduced ATP production and apoptosis *in vitro*, and reduced growth of tumour xenografts *in vivo*. Whilst metabolic defects in WT are not extensively studied, it has been demonstrated that stromal tumours have markedly reduced mitochondrial mass and function compared to blastemal tumours, and that oxidative phosphorylation is considerably lower in WT than normal kidney (48). Taken together with our data, this invokes the possibility that MYCN-regulated over-expression of TOMM20 may alter mitochondrial protein import by the TOM complex and facilitate the glycolytic switch (Warburg effect) in poor prognosis WT. In this regard, it is interesting to note that TOMM20 over-expression has been demonstrated to retard mitochondrial protein import, presumably by disruption of the normal stoichiometry of subunits of the import receptor complex (49).

**Figure 5.**
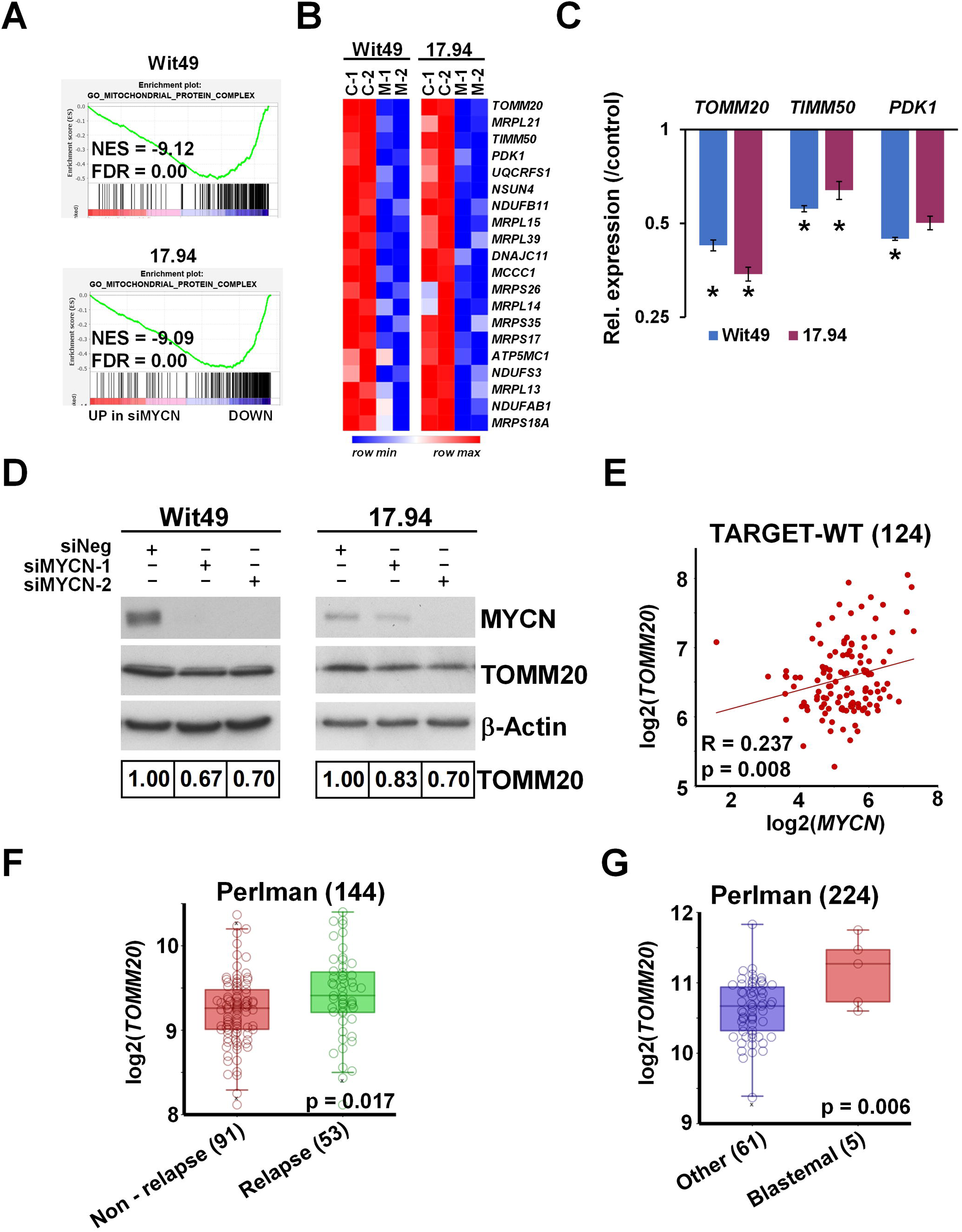
MYCN regulates *TOMM20* and other mitochondrial function genes. **(A)** GSEA showing downregulation of genes encoding for mitochondrial protein complexes with MYCN depletion. **(B)** Heatmap displaying the expression of the top 20 genes in the leading edge of the GO mitochondrial protein complex gene category in MYCN depleted in WT cells. **(C)** Validation of select mitochondrial genes in Wit49 and 17.94 cells by qPCR, 48 hours after MYCN knockdown. A representative of three biological replicates are shown. T-tests were performed on the biological replicates (* p < 0.05). **(D)** Western blot and quantification showing reduction of TOMM20 protein expression after 72 hours of MYCN depletion. (n = 3). **(E)** Expression of *MYCN* and *TOMM20* genes significantly correlated in the TARGET-WT data set (SRP012006). **(F)** Relapsed WT had significantly higher expression of *TOMM20* mRNA than non-relapsed ones, as detected in GSE10320. **(G)** *TOMM20* was also more highly expressed in blastemal WT than in tumours with other histology (GSE31403).

### 3.4 MYCN upregulates methylosome components in WT and influences post-translational regulation via promoting arginine methylation

The methylosome was one of the most enriched GO categories in our transcriptomic analysis of MYCN depletion in WT cells, with six out of the 12 members significantly downregulated in both cell-lines. A heatmap of the methylosome genes shows that the other six members were also downregulated, albeit to a lesser extent (Figure 6A). Expression changes of *WDR77*, encoding for MEP50, and *PRMT1* were validated by using qPCR (Figure 6B). Analysis of RNA-seq data indicated that *PRMT5* was also downregulated in both cell-lines, although to a lesser extent.

**Figure 6.**
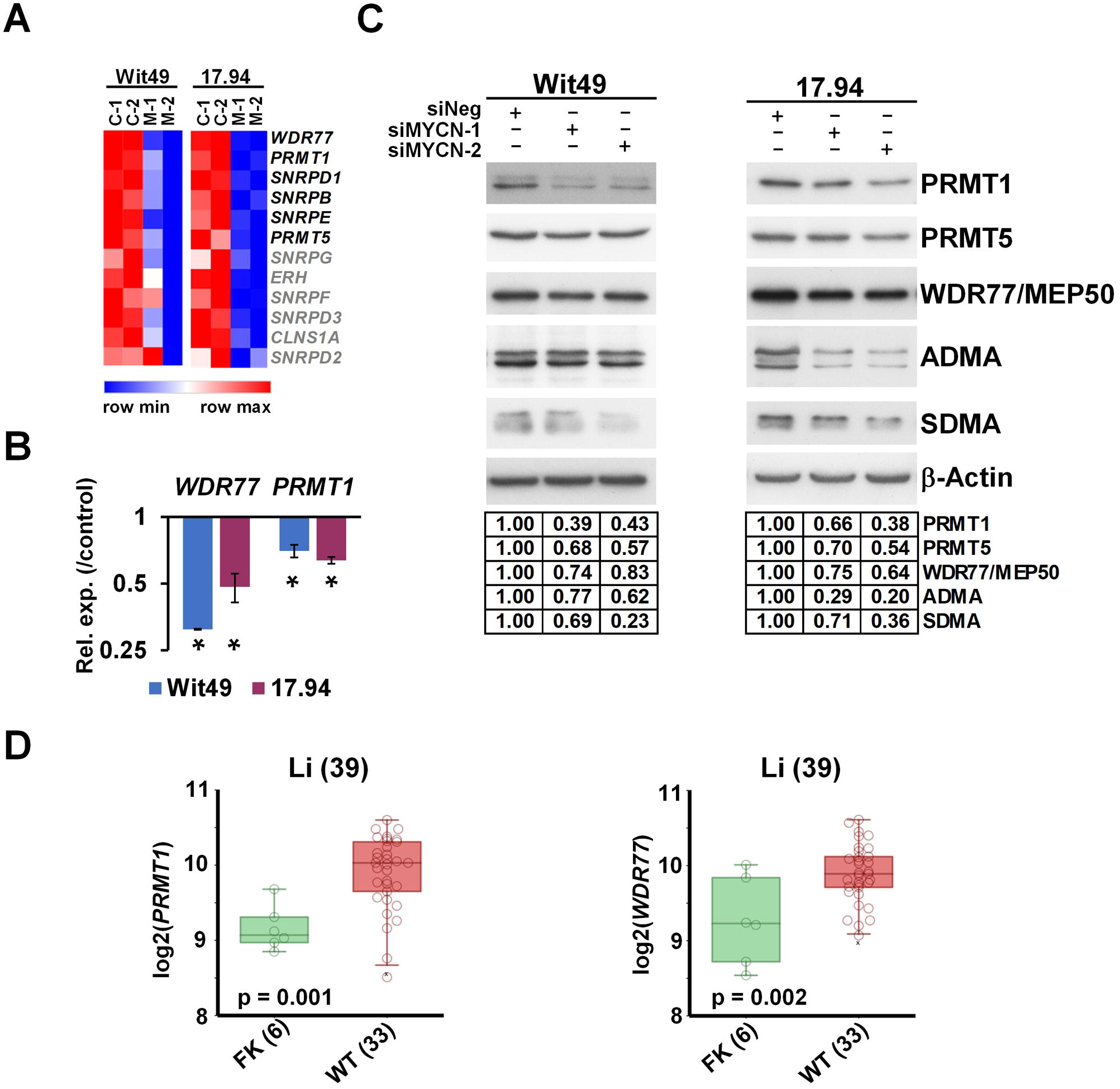
MYCN activates methylosome genes in WT. **(A)** Heatmap of genes encoding for methylosome components, showing expression in MYCN-depleted WT cells. Genes highlighted in black were identified in this study as significantly and substantially regulated by MYCN in two anaplastic WT cells. **(B)** Validation of select methylosome genes in WT cells by qPCR, 48 hours after MYCN knockdown. Statistical significance was calculated based on biological replicates and expression for one representative is shown (* p < 0.05). **(C)** Western blot of select methylosome components in WT cells 72 hours after MYCN knock-down with two different siRNAs (n = 3). Protein expression was calculated based on densitometry and normalized for the loading control on the same filter. Confirmation of MYCN depletion is shown in Figure 5D. **(D)** *PRMT1* and **(E)** *WDR77*, encoding for MEP50, were shown to be expressed at significantly higher levels in WT than in fetal kidney (GSE6120).

Downregulation was confirmed at the protein level for three key methylosome components: PRMT1, catalysing asymmetrical arginine di-methylation (ADMA), PRMT5 and WDR77/MEP50, acting in a complex to effect symmetrical arginine di-methylation (SDMA) modification (Figure 6C). We hypothesized that such profound downregulation of the methylosome components may influence global arginine di-methylation levels, which we tested by using antibodies against ADMA and SDMA modifications. We found that ADMA levels were reduced with MYCN depletion in both cell-lines by using two different siRNAs. SDMA modification was also reduced in the 15–20 kDa size range, suggesting reduction of SDMA marks of snRNP proteins, which participate in RNA splicing.

Our analyses further established that *PRMT1* and *WDR77* were significantly overexpressed in WT as compared to fetal kidney in the transcriptomic data set published by Li et al. (35) (Figure 6D), supporting a role for arginine methyltransferases in WT pathogenesis. Both *PRMT1* and *WDR77* are MYCN targets within the MYCN157 geneset for poor prognosis NB (32) and MYCN also binds the *PRMT5* promoter (50). PRMTs are often over-expressed in cancer (51), and we have previously shown that PRMT5 is a survival factor for MYCN-amplified NB, with PRMT5 interacting with and methylating MYCN protein (52). We have also shown that neuroblastoma cells are sensitive to PRMT1 inhibition (53). Small molecule inhibitors of PRMTs have been developed recently and demonstrated to have efficacy *in vitro* and *in vivo* against cancers such as mantle cell lymphoma (54,55) and are currently in clinical trials for solid tumours and various forms of leukemia (56). Our transcriptomic and protein level analyses suggest that a PRMT-MYCN axis may also be involved in WT, and that selective inhibition of PRMTs may represent a novel targeted therapy for poor prognosis WT.

### 3.5 MYCN represses WT predisposition gene REST, leading to activation of its target genes

Exome sequencing of familial and non-familial Wilms’ tumors recently revealed loss of function mutations in the *REST* gene (encoding RE1 Silencing Transcription Factor) (10). REST is a Krüppel-type zinc-finger transcription factor which acts as a repressor of gene transcription via numerous interactions with chromatin-modifiers, and deregulation of REST is implicated in the pathogenesis of several diseases, including cancer (57). Intriguingly, our RNA-seq revealed *REST* as one of the genes upregulated upon depletion of MYCN. De-repression of REST was also confirmed at the protein level in both cell-lines using two different siRNAs (Figure 7A). Furthermore, we found that the REST-repressed target genes *STMN3, GDAP1* and *ENAH* were decreased after MYCN knockdown, consistent with the upregulation of functional REST protein (Figure 7B). To query the effect of MYCN on REST-regulated genes in WT, we performed GSEA on our MYCN depletion transcriptomic data using gene signatures established in stem cell-derived neurons (58) and embryonic stem cells (ESC) (59) (Figure 7C). Both gene sets were significantly downregulated in both WT cell-lines upon MYCN knockdown, suggesting a MYCN − REST regulatory axis in WT.

**Figure 7.**
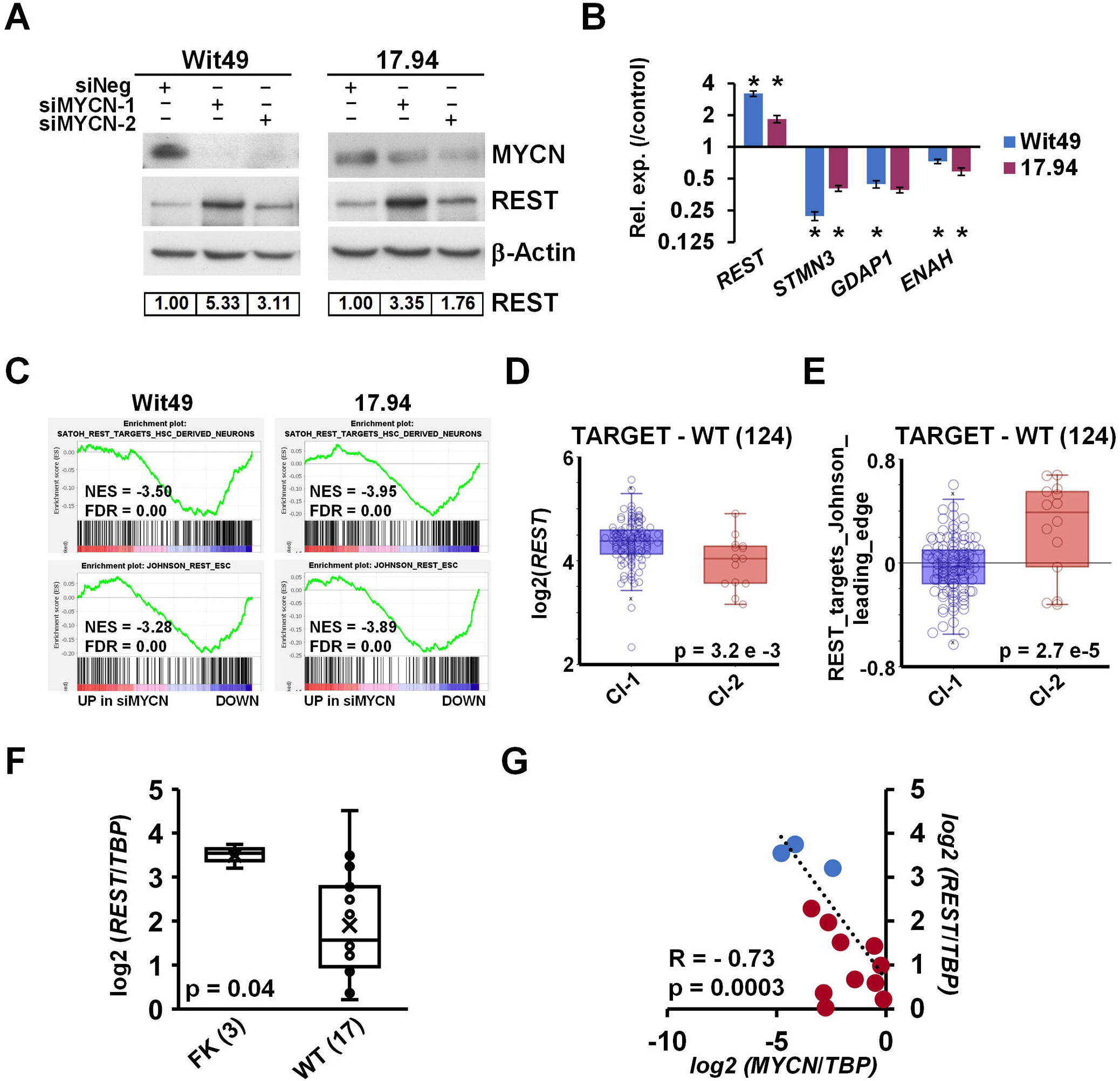
MYCN represses the WT predisposition gene *REST*. **(A)** REST protein is derepressed upon MYCN depletion (n = 3). Protein expression was calculated based on densitometry and normalized for loading control. **(B)** Validation of upregulation of *REST* mRNA and downregulation of its target genes by qPCR after MYCN depletion for 48 hours (n = 3). A representative experiment is shown. T-tests were performed on biological replicates (* p < 0.05). **(C)** GSEA showed downregulation of REST target genes with MYCN depletion. **(D)** REST has a significantly lower expression in cluster 2 tumours in TARGET-WT data set (SRP012006) as compared to cluster 1 (ANOVA). **(E)** REST target genes, showing a significant overall overexpression in cluster 2 WT as compared to cluster 1. (ANOVA) **(F)** *REST* was found to be significantly repressed in WT as compared to FK by qPCR, as assessed by T-test. **(E)** Expression of *MYCN* and *REST* highly and significantly correlated in WT and FK, indicated by red and blue dots, respectively.

*REST* was found to be significantly repressed in cluster 2 of the TARGET – WT data set (Figure 7D), which contained tumours with MYCN signature. In contrast, ESC-specific REST target genes were derepressed in the same group of tumours, as indicated by the expression of REST target metagene, representing overall expression of MYCN regulated REST genes of the signature described by Johnson et al. (59) (Figure 7E). We also found *REST* significantly repressed in WT as compared to fetal kidney in our cohort of primary samples (Figure 7F).

Expression of *MYCN* and *REST* showed a strong, inverse correlation in these tissues (R = −0.73, p = 0.0003) (Figure 7G). Thus our data identifies the repression of the presumptive WT tumour suppressor gene *REST* as a hitherto uncharacterised oncogenic pathway downstream of MYCN deregulation.

Taken together, this study demonstrates that MYCN promotes growth and survival in WT via regulating multiple genes affecting splicing, translation, post-translational modification, microRNAs, metabolism and cellular differentiation (Figure 8). The intersection of MYCN with co-operative oncogenic and tumour suppressor pathways represent possible vulnerabilities of poor-prognosis Wilms’ tumour which can be exploited in the future for urgently required targeted therapeutics.

**Figure 8.**
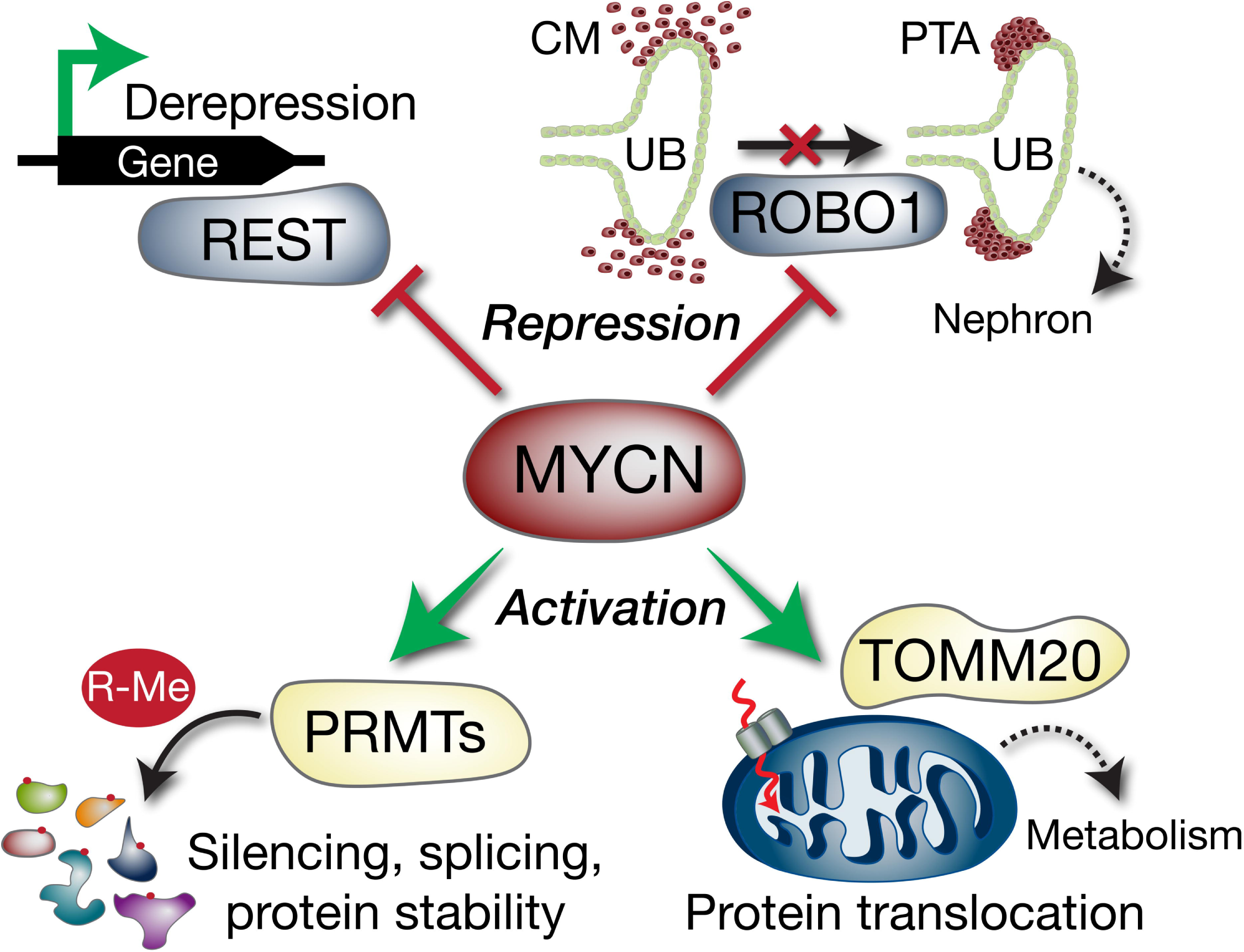
Model depicting select Wilms’ tumour MYCN targets with potential functional consequences. MYCN represses the *REST* tumour suppressor gene, which can result in activation of REST-repressed genes. *ROBO1* is also repressed, which may compromise the differentiation of the condensing mesenchyme (CM) into pre-tubular aggregates (PTA) via inhibition of SLIT-ROBO developmental signalling. The ureteric bud (UB) is also shown. MYCN activation of methylosome genes such as PRMTs alters arginine methylation (R-Me), with subsequent alterations in functions including mRNA splicing and protein stability. Activation of *TOMM20* by MYCN may lead to aberrant mitochondrial protein translocation (dashed red arrow) and altered tumour cell metabolism.

## Legends

**Supplementary figure 1. MYCN immunohistochemistry in adult human kidney**.

MYCN protein is absent in fully differentiated and developed kidney.

**Supplementary figure 2. Cell cycle analysis**.

Depletion of MYCN protein in Wit49 cells by two different siRNAs for 72 hours resulted in a significant increase in the proportion of cells in G2/M phase and a decrease in S phase (n = 3, *** p < 0.01, T-test). No change was observed in the subG1 fraction. MYCN knock-down was confirmed by using Western blot. MYCN knock-down for Wit49 is also shown in Figure 7A.

**Supplementary figure 3. Western blots of MYCN and LIN28B and GSEA of differentiation gene signatures in WT**.

**(A)** Confirmation of MYCN depletion in samples used for RNA-seq. **(B)** Western blot showing downregulation of LIN28B protein after MYCN depletion for 72H in WT cells. Blots of confirmation of MYCN knock-down are shown in Figure 7A for Wit49 and Figure 5D for 17.94. **(C)** GSEA showed upregulation of kidney developmental gene sets in MYCN-depleted WT cells. **(D)** Single cell signatures of differentiated cell types in the fetal kidney were upregulated, while genes associated with proliferating cells were down in MYCN knock-down WT cells.

**Supplementary figure 4. GSEA analysis in transcriptomes of MYCN-depleted WT cells**.

**(A)** Genes overexpressed in WT vs. fetal kidney were mostly downregulated. **(B)** GSEA showed downregulation of MYC/MYCN gene sets. **(C)** GSEA highlighted downregulation of gene sets associated with RNA export from the nucleus and upregulation of unfolded protein response genes.

**Supplementary table 1. Oligonucleotides**

**Supplementary table 2. Antibodies**

**Supplementary table 3. Shared MYCN-regulated genes in Wit49 and 17**.**94**.

**Supplementary table 4. Gene Ontology analysis of MYCN-regulated genes in Wilms’ tumour**.

## Author Contributions

Marianna Szemes: Conceptualization, Methodology, Formal analysis, Investigation, Validation, Data curation, Writing: Original draft, review and editing, Visualization, Supervision. Zsombor Melegh: Formal analysis, Investigation, Validation. Jacob Bellamy, Ji Hyun Park, Biyao Chen: Investigation, Validation. Alexander Greenhough: Visualization. Daniel Catchpoole: Resources. Karim Malik: Conceptualization, Methodology, Formal analysis, Investigation, Resources, Funding acquisition, Supervision, Project administration, Writing: Original draft, review and editing, Visualization.

## Acknowledgments

We wish to thank Prof. Herman Yeger (University of Toronto) and Dr. Keith Brown (University of Bristol) for kindly sharing the WT cell-lines, Wit49 and 17.94, respectively. We also would like to thank Drs. Jane Coghill and Christy Waterfall at the University of Bristol Genomics Facility for assitance with transcriptomics and Dr. Andy Herman for help with flow cytometry. We greatly indebted to Ms Aysen Yuksei and Dr Michael Krivanek for construction of the tissue microarrays and reviewing pathology data.

## Abbreviations

ADMA: asymmetric di-methyl arginine
CM: condensing mesenchyme
DEG: differentially expressed genes
ECL: enhanced chemiluminescence
FACS: fluorescence activated cell sorting
FDR: false discovery rate
FHWT: favourable histology Wilms’ tumour
FK: fetal kidney
GO: GSEA HRP
IHC: Gene ontology
GSEA: Geneset Enrichment Analysis
HRP: horseradish peroxidase
IHC: immunohistochemistry
MNA: MYCN-amplified
NB: neuroblastoma
NES: normalised enrichment score
PTA: pre-tubular aggregates
R-Me: arginine methylation
SDMA: symmetric di-methyl arginine
TIM: translocase of the inner membrane
TMA: tissue microarray
TOM: translocase of the outer membrane
UB: ureteric bud
WT: Wilms’ tumour

