## Supplementary figures and images for "Transcriptomic analyses of MYCN-regulated genes in anaplastic Wilms’ tumour cell lines reveals oncogenic pathways and potential therapeutic vulnerabilities"

### Supplementary figure 1

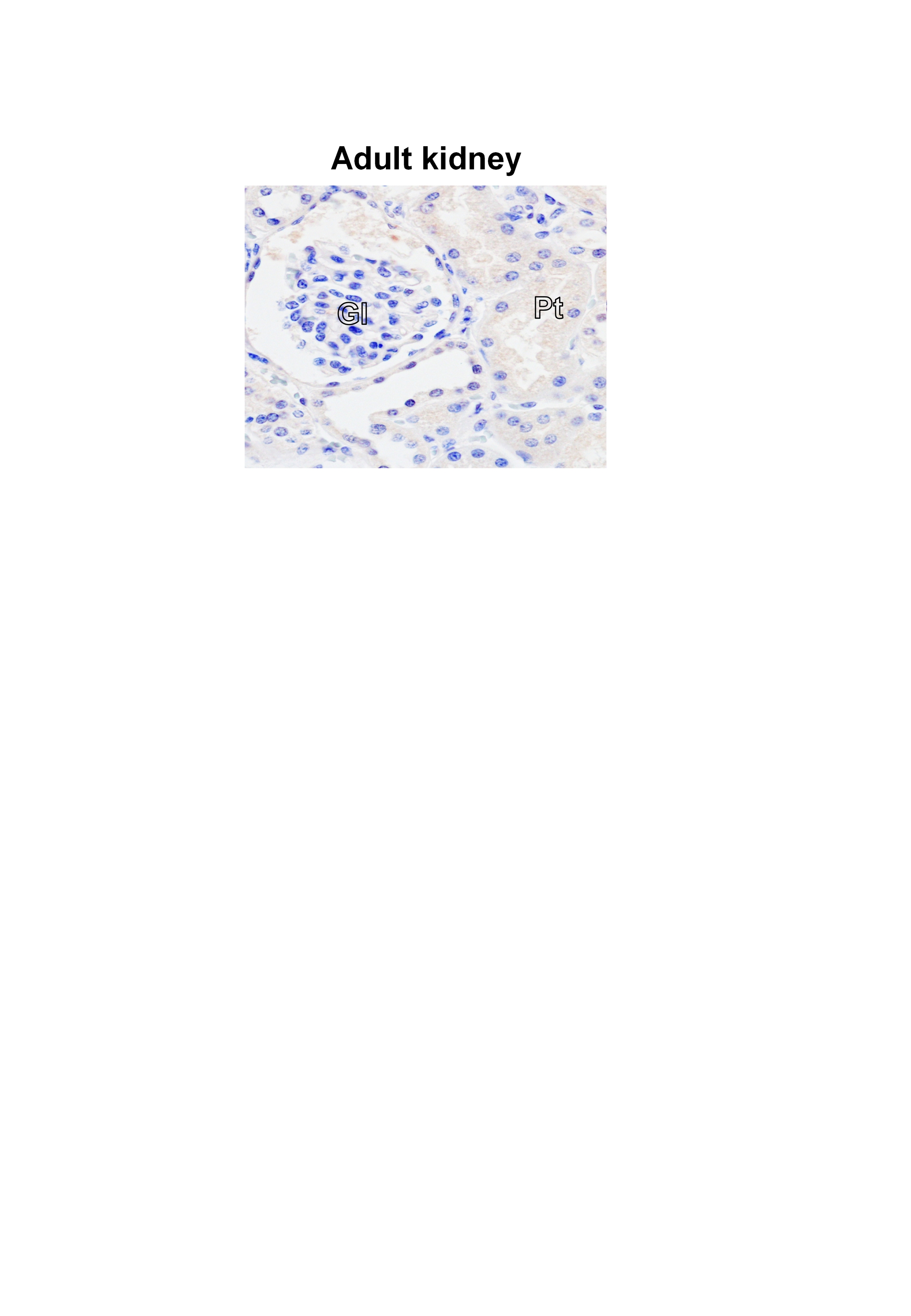

### Supplementary figure 2

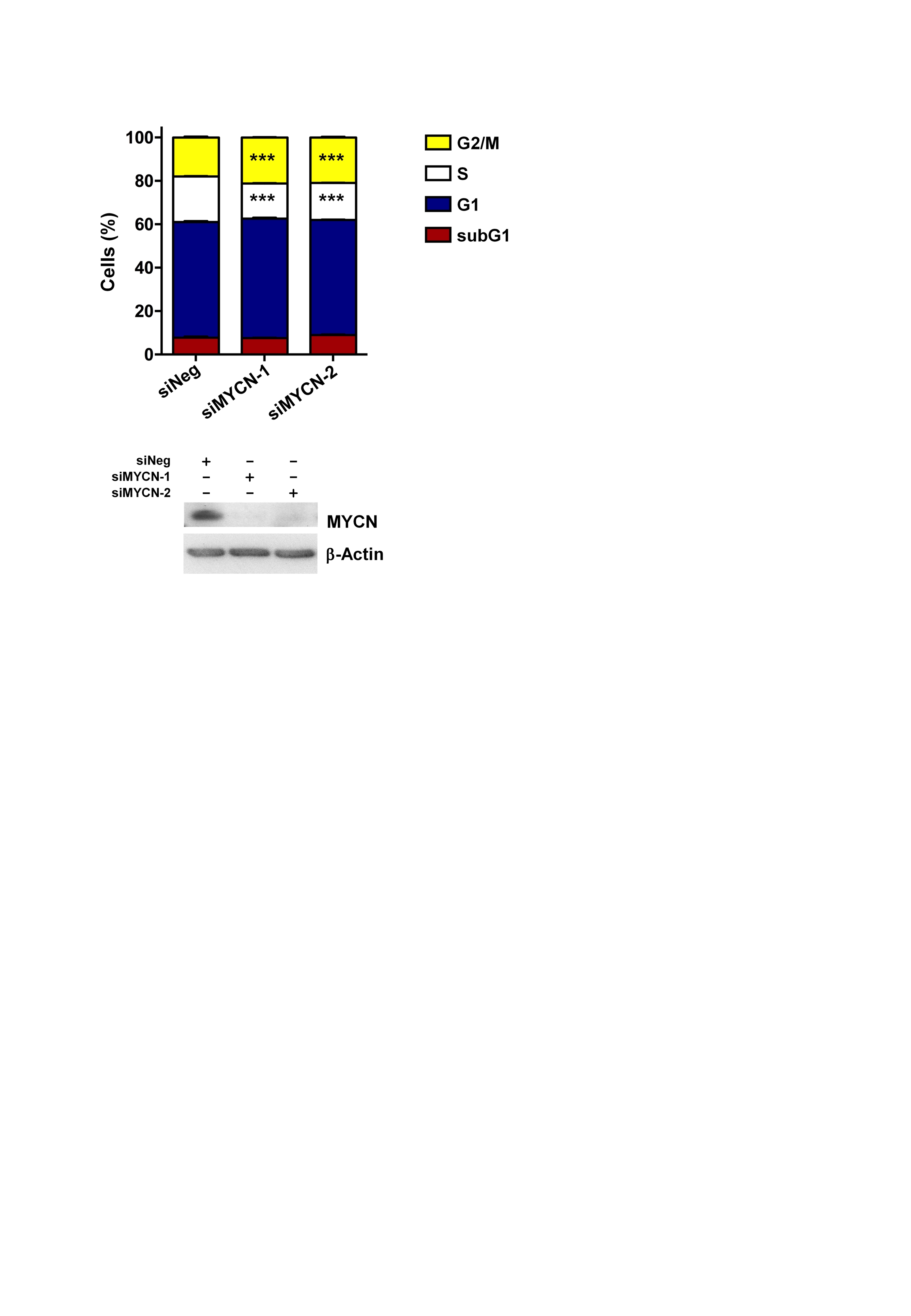

### Supplementary figure 3

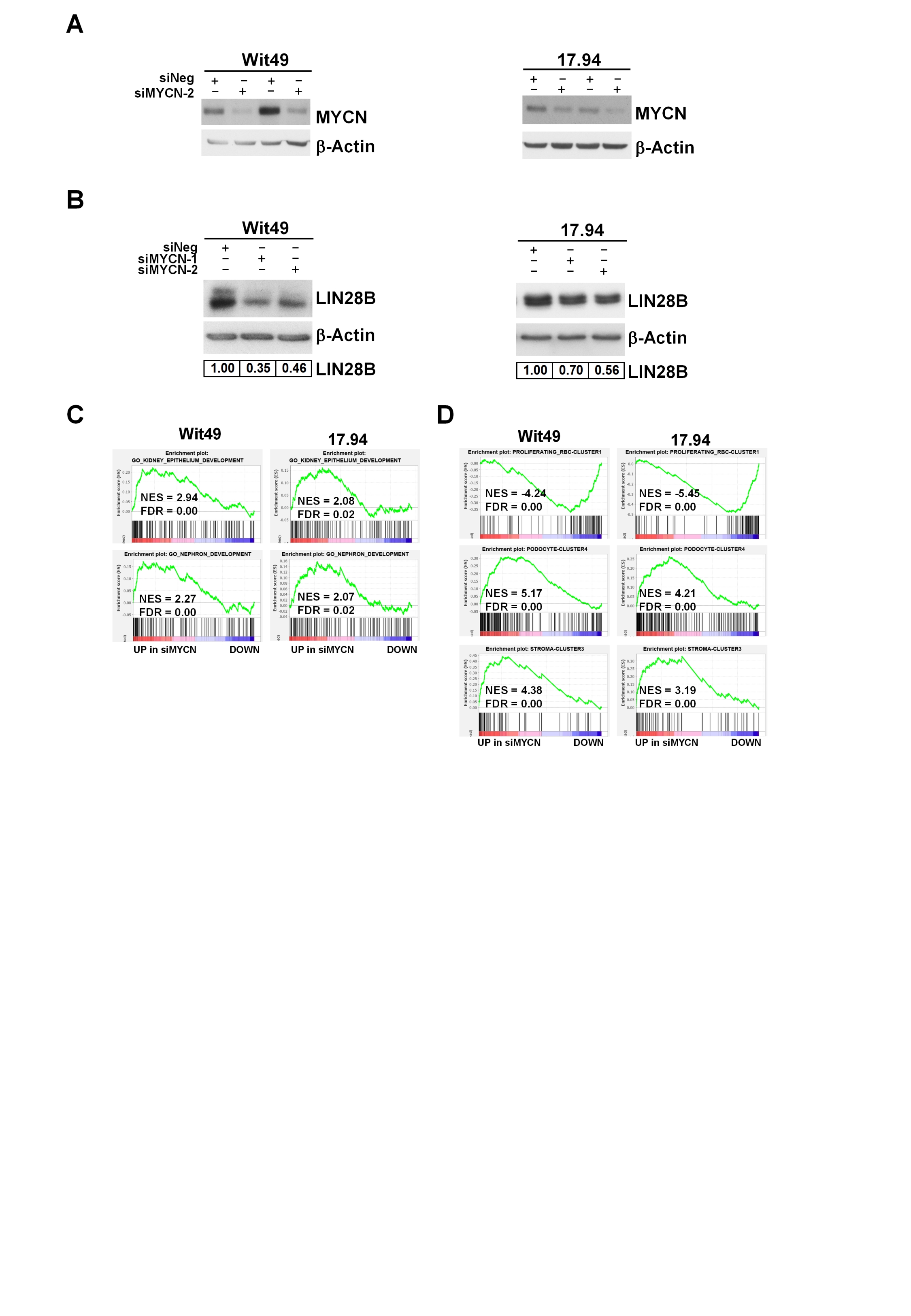

### Supplementary figure 4

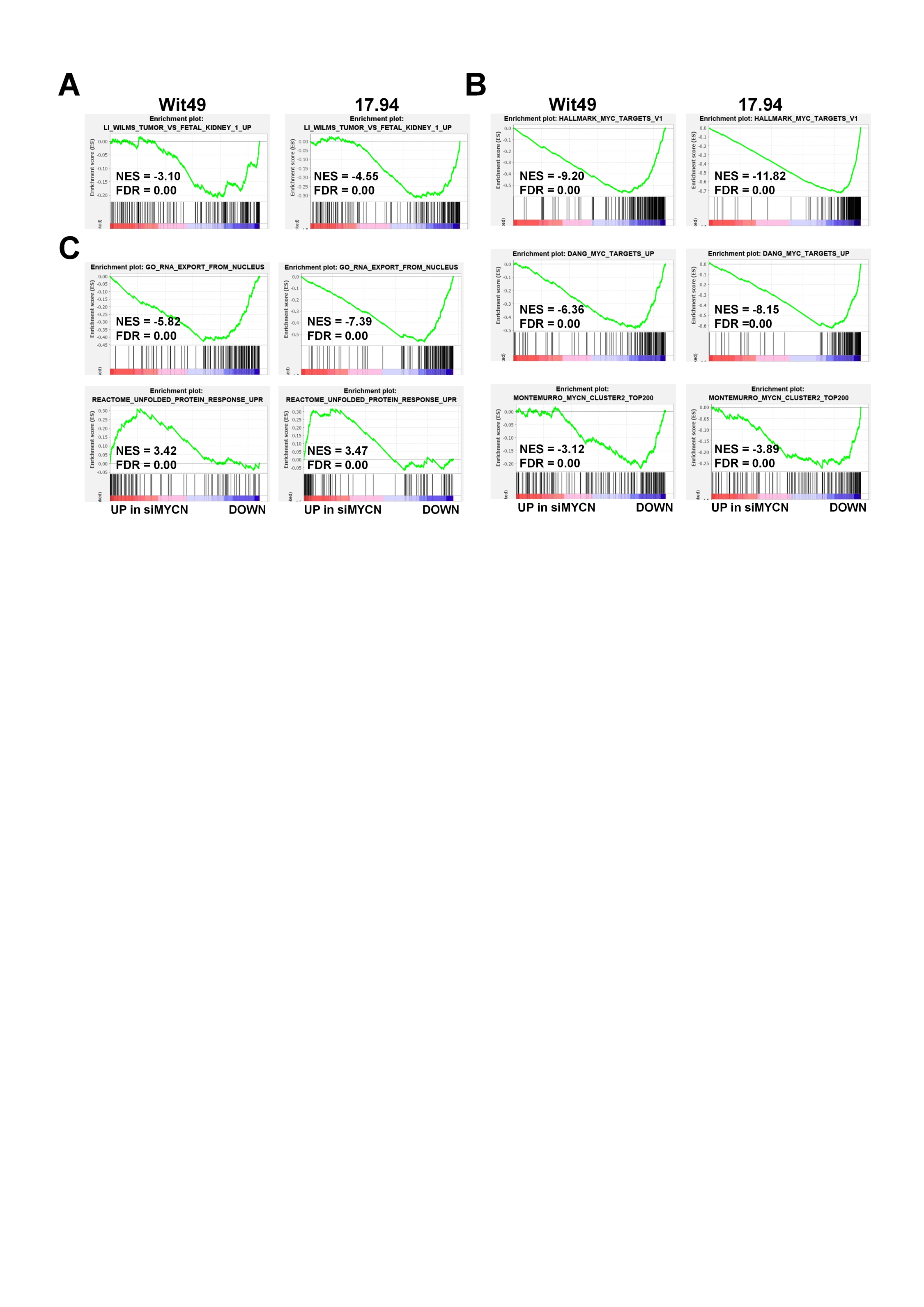
